# Is there a role of phase partitioning in coordinating DNA damage response?

**DOI:** 10.1101/2020.08.26.268763

**Authors:** D. Tosolini, G. Antoniali, E. Dalla, G. Tell

## Abstract

DNA repair pathways are critical processes that need both spatial and temporal fine regulation. Liquid-liquid phase separation (LLPS) is a way to concentrate biochemical reactions, while excluding non-interacting components. Protein’s disordered domains, as well as RNA, favor condensation to modulate this process. Recent insights about phase-separation mechanisms pointed to new fascinating models that could explain how cells could cope with DNA damage responses. In this context, it is emerging that RNA-processing pathways and PARylation events, through the addition of an ADP-ribose moiety to both proteins and DNA, participate in different aspects of the DNA Damage Response (DDR). Remarkably, defects in these regulatory connections are associated with genomic instability and human pathologies. In addition, it has been recently noticed that several DNA repair enzymes, such as 53BP1 and APE1, are endowed with RNA binding abilities. APE1 is a multifunctional protein belonging to the Base Excision Repair (BER) pathway of non-distorting DNA lesions, bearing additional ‘non-canonical’ DNA-repair functions associated with processes coping with RNA metabolism. In this work, after reviewing the recent literature supporting a role of LLPS in DDR, we analyze, as a proof of principle, the interactome of APE1 using a bioinformatics approach to look for clues of LLPS in BER. Some of the APE1 interactors are associated with cellular processes in which LLPS has been either proved or proposed and are involved in several tumorigenic and amyloidogenic events. This work represents a paradigmatical pipeline for evaluating the relevance of LLPS in DDR.

**Statement of significance:** In this work, we aimed to test the hypothesis of an involvement of phase-separation in regulating the molecular mechanisms of the multifunctional enzyme APE1 starting from the analysis of its recently-characterized protein-protein interactome (PPI). We compared APE1-PPI to phase-separation databases and we performed functional enrichment analysis, uncovering links between APE1 and already known demixing factors, establishing an association with liquidliquid phase separation. This analysis could represent a starting point for implementing downstream experimental validations, using in vitro and in vivo approaches, to assess actual demixing.

## Phase separation in nuclear organization and functions related to DNA damage response

Nuclear dynamics, among other crucial cellular processes, has been recently established to be tuned, at least partly, through the widespread phenomenon of phase separation (1). After a decade of active research, it is now accepted that the demixing process is a thermodynamically-driven phenomenon, giving rise to a variety of dynamic bodies primarily composed of nucleic acids and proteins (2), interacting through quinary interactions (3) mostly involving unstructured portions of proteins, especially intrinsically disordered regions (IDRs)(4, 5). Some examples of nuclear processes proposed to be shaped by phase separation are: heterochromatin domains formation, transcription, nucleolar metabolism and DNA damage response (DDR). Indeed, it has been shown that chromatin structure dynamics may be regulated through phase-separation of several proteins (e.g. HP-1 and BRD4) involved in the reading of epigenetic marks on histone tails (6–8); histones tail-DNA interactions, as well, might have a role in this process (8). Moreover, an outstanding example of phase separation in the nucleus is represented by the nucleolus, the cellular body devoted to ribosome biogenesis. In recent years, it has been demonstrated that nucleoli arise by phase separation induced by transcription of rRNAs from their genomic loci (9, 10). Strikingly, the nucleolar ultrastructure, composed of three domains arranged in a shell-core manner, represents a proof of concept of multiple, nested demixed liquid compartments mirroring the different functions of the three layers (11). Another process for which a phase separation mechanism has been proposed is represented by DNA transcription: recent researches shed new light onto the actual mechanism of recruitment of transcription factors, proposing cooperative kinetics to explain the effects driven by enhancers and superenhancers, via demixing processes of transcription factors themselves (12, 13). However, transcription, as well as other processes claimed to be driven by phase separation (e.g. heterochromatinization), remains to be fully characterised, because it differs from biomolecular condensates in some aspects, which are reviewed in (1). In particular, some compartments characterized by many phase-separating features do not respect the canonical features of liquid biocondensates (namely the round shape, no shear elasticity and internal dynamics), raising the question of whether phase-separation could be displayed in several different aspects. For example, paraspeckles, although regarded as demixed bodies upon NEAT1 increase, show a one-axis preferential growth, unusual for LLPS-based granules. Heterochromatic domains, instead, which were proposed to form by phase separation because of their apparent properties of coalescing, for excluding inert probes and for causing density transition in HP-1 distribution, were finally shown to undergo their round shape degeneration several cell cycles after phase separation occurrence, thus denying the initial hypothesis (14).

## Role of RNA transcription and PARylation in promoting recruitment of DNA repair enzymes on the site of the lesion (examples from DSBs- and SSBs-repair)

RNA is a crucial element of phase-demixed compartments, and some studies shown its involvement in DNA repair. A first evidence of this involvement was given by the DDR-related action of retrotransposons in yeast: indeed, it was shown that retrotransposon elements might replace homologous sequences and become integrated at the lesion site (15, 16). In (17), the authors suggested another link existing between retrotransposons and DDR: while reverse transcriptases could promote repair by canonical transcripts, integrases might promote cDNA insertion and cDNA might act as template to bridge the DSB, leading to repair by “*in trans*” or “*in cis*” mechanisms. Additionally, with regard to repair of DSBs, it was recently suggested that Rad52 might promote transcript-dependent DSB repair through inverse strand exchange, likely followed by reverse transcription of a ssDNA overhang (18). Rad52 has also been demonstrated to lead to transcript-dependent DSB repair (19, 20). This perspective is supported by the mounting evidence accounting for R-loops as physiological regulators in the genome (21, 22). Notably, Rad52 (in yeast) and FUS (in human) were observed to contribute to the formation of a molecular biocondensate at DSB sites, carrying out different roles, namely the organization of nuclear microtubule filaments (23), protecting the resected end of lesions and promoting DNA-damage signaling (24, 25), as well as recruiting other DSB-repair-related enzymes (26). Analogously, 53BP1, which is known to be important for DSB signaling and to affect the progression of cell cycle, was demonstrated to take part in liquid compartments, too (24, 25). Recently, it was found that ncRNAs seem to have a critical role in the formation of a liquid compartment at the DSB site. In detail, a novel class of RNAs has been defined and named DDRNAs: they are produced from the processing of dilncRNAs (damage-induced long non coding RNAs), which are transcribed at DSBs foci in a bidirectional manner (27, 28). DDRNAs are guided to the lesion site by dilncRNAs and both of them are supposed to contribute to the recruitment of repair enzymes (27, 29).

A particular example showing the involvement of RNA in DDR is PARylation: this is a reversible post-translational modification leading to the addition of ADP-ribose molecules to both proteins and DNA, it is carried out by the PARP enzymes family and it represents one of the principal signals of genomic damage in cells (30, 31). Interestingly, PARylation, that might be tightly related to phase separation due to the role of RNA as a key scaffolding element of such compartments (32), has been found to direct the assembly of FUS upon DNA repair sites (33) and to be essential, along with DNA, for SSB (Single-strand breaks) repair associated with XRCC1 (34).

Since PARylation seems to trigger DDR pathways through phase separation (35), it could be speculated that PARP-1 might play a similar role in the regulation of BER pathway functions, through recognition of SSBs. Indeed, PARP-1 was shown to interact with several BER factors (e.g. XRCC1, POLβ and LIG3) and to modulate the activity of glycosylases (36) and the 3’-exonuclease activity of APE1 (37). Work is needed along these lines to prove this hypothesis, which could explain why PARP-1 inhibition significantly hampers the efficiency of the BER pathway (38).

## Interactomes of DNA repair enzymes: focus on RNA processing proteins

Along with the involvement of RNA molecules themselves, it is widely accepted that the interaction with RNA represents a key feature of most DDR enzymes, both in direct and indirect manners. It was shown that many RNA binding proteins (RBPs) are required to ensure proper production of DDR factors (as reviewed in (39)), thus indirectly influencing the repair process. Nonetheless, RBPs were shown to directly take part in DDR, as enzymes involved in mRNA and miRNA processing have been associated with DNA repair, as well. For instance, RBM14 is a RBP involved in alternative splicing and it is recruited to DSB sites via PARP1 (40, 41); likewise, HNRNPD is necessary for the DNA resection step in the homologous recombination pathway (42). Helicases, like for example DEAD-box helicases, are interesting RNA-interacting proteins involved in RNA metabolism (43, 44) and in DDR (45, 46) and some of them were shown to take part in demixing bodies (47, 48).

Interestingly, the small non-coding RNA machinery including DICER and DROSHA are important for DSB repair and might possibly have a role in the establishment of the inverse strand exchange (28, 49).

## Bioinformatics analysis of APE1 and demixing proteins: analysis of the APE1 interactome suggests a novel hypothesis for triggering of Base Excision Repair pathway

Based on the above-mentioned literature insights, the Apurinic/Apyrimidinic Endonuclease APE1 looks like an appealing subject for investigating its interactome and its participation in demixing processes. APE1 is a fundamental BER enzyme, acting as the main AP-endonuclease in mammalians, with many recently characterized non-canonical functions, particularly in RNA metabolism, including taking part in the biogenesis of ncRNAs, possibly through the interaction with DROSHA (50) or with other phase separating factors (e.g. NPM-1) (51), as well as with several RNA species (52). These novel functions might be explained by hypothesizing a phase separation mechanism for APE1 recruitment.

Here, we aimed to give a proof of principle on how bioinformatic resources might be employed to gain some useful insights on the LLPS world, defining a pipeline to help directing the following experimental activity. In our recent work (53), we defined an APE1 interactome that we here employed in the following analysis pipeline to look for clues of LLPS in BER. In the first step, we retrieved the disorder content of APE1 protein-protein interactors (APE1-PPI) from the MobiDB database (version 3.1.0) (54), integrating manual reviews and *in silico* predictions. Out of 515 interactors, we were able to retrieve the entries for about 350 proteins and examined the distribution of these values. Most of the interactors were characterized by a low disorder content (below 20%) (Fig. 1).

**FIGURE 1.**
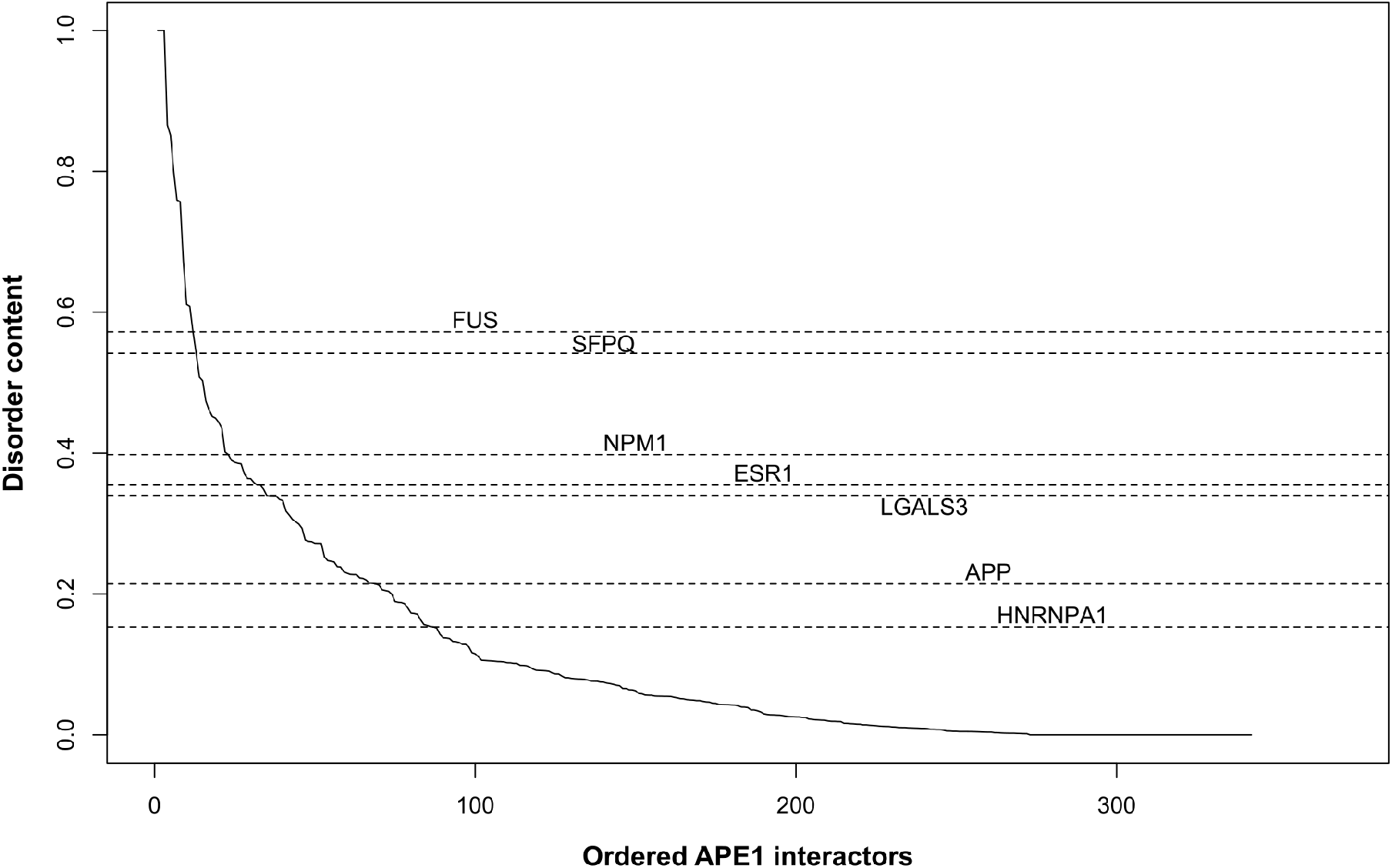
Distribution of the disorder content of the APE1 interactome proteins. The interactors, ordered by decreasing disorder content as retrieved from MobiDB, are identified by their position in the list of interactors.

Then, we intersected the APE-PPI list with data from PhaSePro (55), a manually curated collection of proteins characterised as demixing *in vivo*. The output consisted of seven interactors, namely APP, NPM1, LGALS3, HNRNPA1, FUS, SFPQ and ESR1, whose disordered content was compared to the general distribution (not shown).

We noticed that all the reviewed demixing proteins were characterized by an internal disorder content greater than 0.15; thus, we focused on those PPI having a disordered content above that threshold, defining a subset made of 88 members. We compared them to entries in PhaSepDB (56), which aggregates a wide range of direct and indirect evidence of proteins phase separation (e.g. fully demonstrated or suggested by high-throughput data), reaching a final set of 49 likely demixing interactors (Table S1 in the Supporting Material).

To gain some insights on the biological processes involving these proteins, we performed a functional enrichment analysis (57) employing ClueGO (58), a Cytoscape (59) plugin allowing to use different ontologies/databases, focusing on biological processes (Fig. 2A) and intracellular localization (Fig. 2B).

**FIGURE 2.**
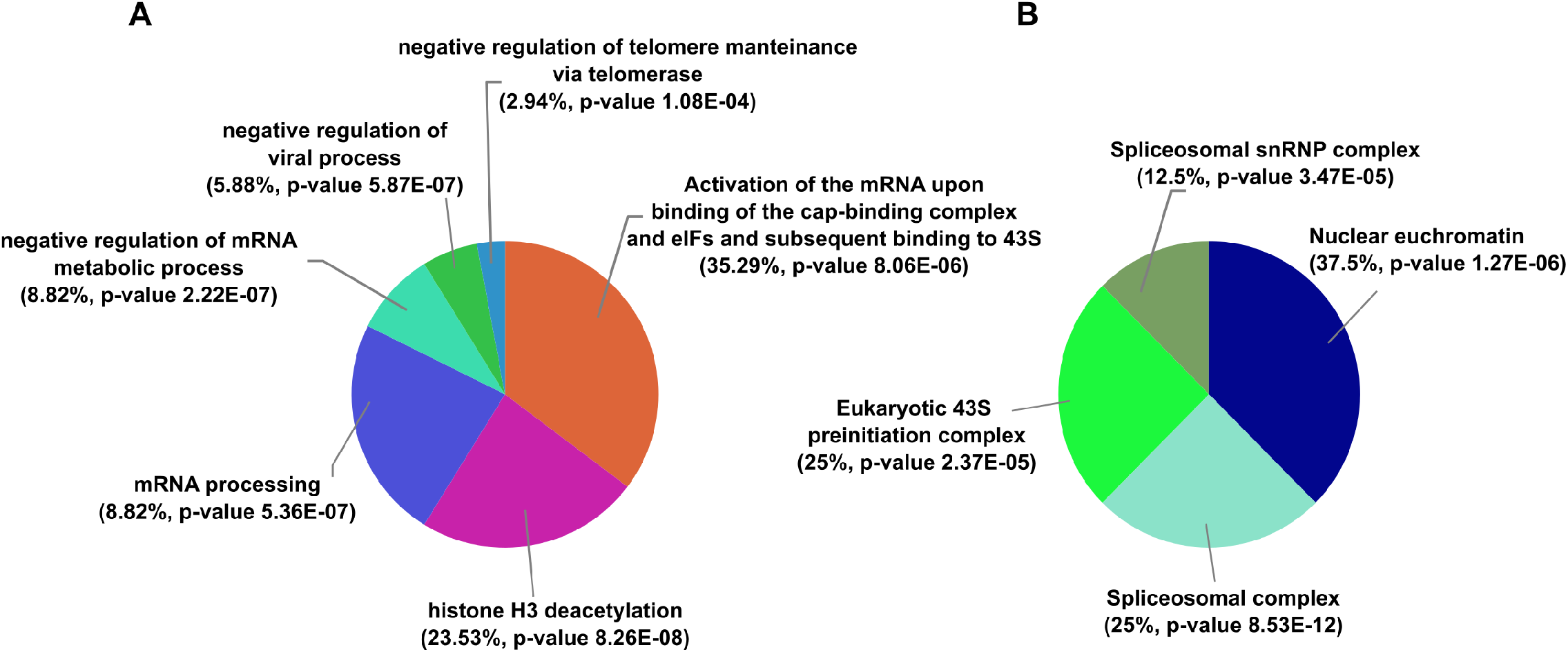
Functional enrichment analysis of the set of 49 interactors likely correlated to phase separation. (**A**) Enriched biological processes. (**B**) Enriched cellular compartments. Percent values refer to the amount of enriched terms associated with each cluster.

We first analysed the set of 49 likely demixing interactors defined by PhaSepDB, and the results highlighted 34 significantly enriched terms, furtherly associated to six major processes by ClueGO, as shown in Fig. 2A (detailed list of enriched terms in Fig. S1). Interestingly, more than half of these terms were associated with gene expression and RNA processing, while the remaining terms were associated with viral and telomeric regulation.

A second analysis on the same gene set took into consideration the intracellular localization of the above mentioned PPI. From this analysis, enriched terms related to euchromatin, spliceosome and translation preinitiation complex, all of which are cellular departments strictly linked to the metabolism of nucleic acids, which might act as dynamic scaffolds for liquid-like structures (Fig. 2B).

We performed the same analysis on the set of seven PhaSePro, fully characterized interactors. This investigation did not yield any significant results (not shown); for this reason, we added APE1 to slightly increase the complexity of the examined dataset, identifying three functional areas (Fig. 3). Interestingly, one of the enriched terms pointed to the formation of amyloids, which are a common hallmark of neurodegenerative diseases, often related to liquid-demixing proteins (60, 61).

**FIGURE 3.**
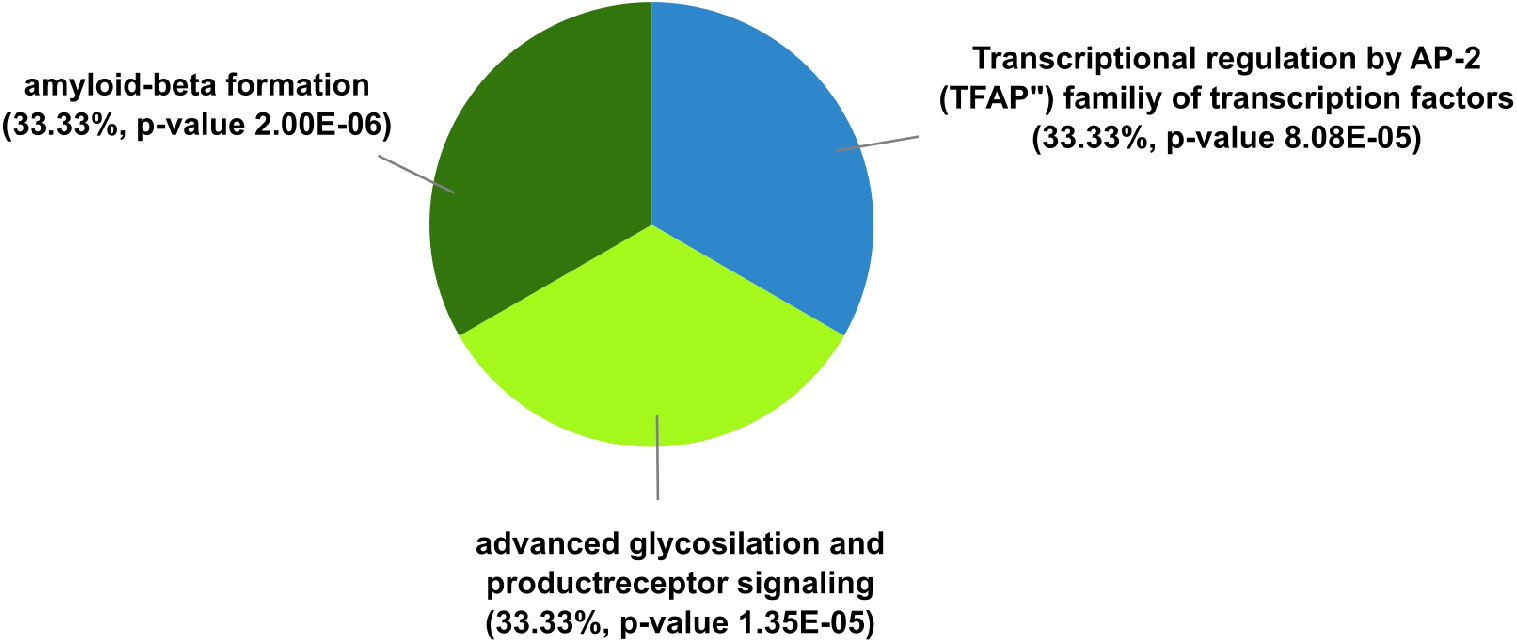
Functional enrichment analysis of the extended set of PhaSePro characterized proteins (including APE1). Enriched biological processes interestingly suggest a role in amyloids formation, regulation of transcription and AGE receptor signalling.

Lastly, we compared the list of 49 interactors to the MSigDB database, a collection of gene sets co-expressed and/or involved in physiological and pathological processes (i.e. molecular signatures) that functionally complements the ontologies previously investigated using ClueGO (62, 63). We considered several gene sets, taking into account the first 50 enriched results. We obtained a significant association with different kind of cancer signatures, namely liver, prostate and haematological tumors; in addition, signatures were also linked to the activation of the PBMCs (i.e. peripheral blood mononucleated cells). Finally, enriched terms also pointed to ribogenesis and mRNA splicing, consistently with previous ClueGO results: these terms strongly suggest a relation, both physical and functional, with nucleic acids (especially RNA), with their metabolism and with compartments in which they accumulate (Fig. S2 and Table S2). We hypothesize that this connection might happen through a direct or indirect recruitment to liquid demixing bodies, the impairment of which could be related to cancerous conditions. This particular association with nucleic acids might also represent an evidence for a novel APE1 function related to phase-separation: on such basis, more investigation is demanded to elucidate the possible role of biocondensates in BER pathway triggering, explaining the role of these RNA-interacting partners, and also for establishing novel protocols for drug development.

## Conclusive remarks

This work describes a bioinformatics pipeline to pinpoint new phase separating proteins candidates, starting from experimental datasets of protein interactomics data, although the underlying direct or indirect molecular mechanisms are still to be demonstrated. We are aware that in (64) the Authors suggested a similar approach employing an ontology terms enrichment analysis; our perspective, however, is substantially different, since we performed the analysis on the interactome of a single protein, with details about phase separation that remain still elusive. The results obtained through this bioinformatics pipeline will be useful in later experimental validation, to identify demixing cofactors that might unveil to be crucial in reproducing phase-separation in vitro (65). For example, on the basis of the close relationship between APE1 and NPM1, the natural follow up of this investigation could be represented by exploring the possible joint phase-separation of these proteins in the presence of rRNA, which is known to be required for NPM1 phase separation and that might also be necessary to APE1 demixing. Besides, drugs selectively impairing LLPS might be employed to demonstrate such phenomenon: 1,6-hexanediol, for example, was used in previous works to show the demixed status of some bodies. Some doubts have been raised about this proving technique, since this molecule might produce artifacts in living cells; besides, its amphipathic nature is not able to impair some of the most common molecular interactions giving rise to BMCs. Moreover, we suggest that the approach we showed here might give a contribution in uncovering new molecular strategies for the therapy of human diseases that have been recently linked to phase partitioning, especially neurodegenerative diseases (e.g. Alzheimer disease, amyotrophic lateral sclerosis (ALS), frontotemporal dementia). In fact, well known ALS-related mutations, such as substitutions in the disordered C-terminus of TDP-43 and hexanucletide expansion in C9orf72, were shown to impair phase separation of these proteins, suggesting their relevance in the onset of the pathological condition (66, 67). Additionally, altered proteostasis, which leads to the formation of aggregates in neuronal cells, is a hallmark of such conditions and was related to phase separation. Nonetheless, a full understanding of how this aggregation influences the pathological outcome is missing and needs deeper investigations to elucidate the exact relationship linking the fiber formation and the toxic effect (68). Additionally, an interesting question arising from the hypothesis of pathological misregulation of liquid compartments concerns how these bodies are physiologically maintained and what are the impaired regulatory mechanisms leading to unreversible aggregation, if any (68). Lastly, since anti-cancer therapies are partly based on inefficient DNA repair, further characterization of molecular mechanisms and associated dynamics, along with advanced unfolding of the interactomes involved in such pathways, might uncover new oncological targets. Therefore, a better understanding of the mechanisms underlying LLPS will certainly improve our knowledge on how to target deregulated processes of the DDR, selectively impacting on several human pathologies.

## Supporting Material

Details of Material and Methods and supplementary figures and tables are available at XXY.

## Author contributions

GT designed the paper outline and the research plan. DT performed the analysis. GA and ED analyzed data. DT, GA and ED wrote the manuscript. GT supervised the writing of the whole Manuscript.

## Acknowledgments

This work was supported by a grant from Associazione Italiana per la Ricerca sul Cancro (AIRC) (IG19862) to G. Tell.

## References

1. Peng, A., and S.C. Weber. 2019. Evidence for and against liquid-liquid phase separation in the nucleus. Non-coding RNA. 5.

2. Banani, S.F., H.O. Lee, A.A. Hyman, and M.K. Rosen. 2017. Biomolecular condensates: Organizers of cellular biochemistry. Nat. Rev. Mol. Cell Biol. 18:285–298.

3. Ribeiro, S., S. Ebbinghaus, and J.C. Marcos. 2018. Protein folding and quinary interactions: creating cellular organisation through functional disorder. FEBS Lett. 592:3040–3053.

4. Protter, D.S.W., B.S. Rao, B. Van Treeck, Y. Lin, L. Mizoue, M.K. Rosen, and R. Parker. 2018. Intrinsically Disordered Regions Can Contribute Promiscuous Interactions to RNP Granule Assembly. Cell Rep. 22:1401–1412.

5. Lin, Y., D.S.W. Protter, M.K. Rosen, and R. Parker. 2015. Formation and Maturation of Phase-Separated Liquid Droplets by RNA-Binding Proteins. Mol. Cell. 60:208–219.

6. Erdel, F., and K. Rippe. 2018. Formation of Chromatin Subcompartments by Phase Separation. Biophys. J. 114:2262–2270.

7. Sanulli, S., M.J. Trnka, V. Dharmarajan, R.W. Tibble, B.D. Pascal, A.L. Burlingame, P.R. Griffin, J.D. Gross, and G.J. Narlikar. 2019. HP1 reshapes nucleosome core to promote phase separation of heterochromatin. Nature. 575.

8. Gibson, B.A., L.K. Doolittle, M.W.G. Schneider, L.E. Jensen, N. Gamarra, L. Henry, D.W. Gerlich, S. Redding, and M.K. Rosen. 2019. Organization of Chromatin by Intrinsic and Regulated Phase Separation. Cell. 179:470–484.e21.

9. Smirnov, E., J. Borkovec, L. Kováčik, S. Svidenská, A. Schröfel, M. Skalníková, Z. Švindrych, P. Křížek, M. Ovesný, G.M. Hagen, P. Juda, K. Michalová, M. V. Cardoso, D. Cmarko, and I. Raška. 2014. Separation of replication and transcription domains in nucleoli. J. Struct. Biol. 188:259–266.

10. Falahati, H., B. Pelham-Webb, S. Blythe, and E. Wieschaus. 2016. Nucleation by rRNA Dictates the Precision of Nucleolus Assembly. Curr. Biol. 26:277–285.

11. Feric, M., N. Vaidya, T.S. Harmon, D.M. Mitrea, L. Zhu, T.M. Richardson, R.W. Kriwacki, R. V. Pappu, and C.P. Brangwynne. 2016. Coexisting Liquid Phases Underlie Nucleolar Subcompartments. Cell. 165:1686–1697.

12. Sabari, B.R., A. Dall’Agnese, A. Boija, I.A. Klein, E.L. Coffey, K. Shrinivas, B.J. Abraham, N.M. Hannett, A. V. Zamudio, J.C. Manteiga, C.H. Li, Y.E. Guo, D.S. Day, J. Schuijers, E. Vasile, S. Malik, D. Hnisz, T.I. Lee, I.I. Cisse, R.G. Roeder, P.A. Sharp, A.K. Chakraborty, and R.A. Young. 2018. Coactivator condensation at super-enhancers links phase separation and gene control. Science (80-.). 361:eaar3958.

13. Nair, S.J., L. Yang, D. Meluzzi, S. Oh, F. Yang, M.J. Friedman, S. Wang, T. Suter, I. Alshareedah, A. Gamliel, Q. Ma, J. Zhang, Y. Hu, Y. Tan, K.A. Ohgi, R.S. Jayani, P.R. Banerjee, A.K. Aggarwal, and M.G. Rosenfeld. 2019. Phase separation of ligand-activated enhancers licenses cooperative chromosomal enhancer assembly. Nat. Struct. Mol. Biol. 26:193–203.

14. Strom, A.R., A. V. Emelyanov, M. Mir, D. V. Fyodorov, X. Darzacq, and G.H. Karpen. 2017. Phase separation drives heterochromatin domain formation. Nature. 547:241–245.

15. Moore, J.K., and J.E. Haber. 1996. Capture of retrotransposon DNA at the sites of chromosomal double-strand breaks. Nature. 383:644–646.

16. Morrish, T.A., N. Gilbert, J.S. Myers, B.J. Vincent, T.D. Stamato, G.E. Taccioli, M.A. Batzer, and J. V. Moran. 2002. DNA repair mediated by endonuclease-independent LINE-1 retrotransposition. Nat. Genet. 31:159–165.

17. Meers, C., H. Keskin, and F. Storici. 2016. DNA repair by RNA: Templated, or not templated, that is the question. DNA Repair (Amst). 44:17–21.

18. McDevitt, S., T. Rusanov, T. Kent, G. Chandramouly, and R.T. Pomerantz. 2018. How RNA transcripts coordinate DNA recombination and repair. Nat. Commun. 9:1–10.

19. Storici, F., K. Bebenek, T.A. Kunkel, D.A. Gordenin, and M.A. Resnick. 2007. RNA-templated DNA repair. Nature. 447:338–341.

20. Keskin, H., Y. Shen, F. Huang, M. Patel, T. Yang, K. Ashley, A. V. Mazin, and F. Storici. 2014. Transcript-RNA-templated DNA recombination and repair. Nature. 515:436–439.

21. Niehrs, C., and B. Luke. 2020. Regulatory R-loops as facilitators of gene expression and genome stability. Nat. Rev. Mol. Cell Biol. 21:167–178.

22. Tan, J., M. Duan, T. Yadav, L. Phoon, X. Wang, J.M. Zhang, L. Zou, and L. Lan. 2020. An R-loop-initiated CSB-RAD52-POLD3 pathway suppresses ROS-induced telomeric DNA breaks. Nucleic Acids Res. 48:1285–1300.

23. Oshidari, R., R. Huang, M. Medghalchi, E.Y.W. Tse, N. Ashgriz, H.O. Lee, H. Wyatt, and K. Mekhail. 2020. DNA repair by Rad52 liquid droplets. Nat. Commun. 11.

24. Kilic, S., A. Lezaja, M. Gatti, E. Bianco, J. Michelena, R. Imhof, and M. Altmeyer. 2019. Phase separation of 53 BP1 determines liquid-like behavior of DNA repair compartments. EMBO J. 38:1–17.

25. Mirza-Aghazadeh-Attari, M., A. Mohammadzadeh, B. Yousefi, A. Mihanfar, A. Karimian, and M. Majidinia. 2019. 53BP1: A key player of DNA damage response with critical functions in cancer. DNA Repair (Amst). 73:110–119.

26. Lenken, S.C., B.R. Levone, G. Filosa, M. Antonaci, F. Conte, C. Kizilimark, S. Reber, A. Loffreda, F. Biella, A.E. Ronchi, O. Muhlemann, A. Bachi, M.-D. Ruepp, and S.M.L. Barabino. 2019. FUS-dependent phase separation initiates double-strand break repair. bioRxiv. (preprint posted Oct. 11th, 2019).

27. Pessina, F., F. Giavazzi, Y. Yin, U. Gioia, V. Vitelli, A. Galbiati, S. Barozzi, M. Garre, A. Oldani, A. Flaus, R. Cerbino, D. Parazzoli, E. Rothenberg, and F. d’Adda di Fagagna. 2019. Functional transcription promoters at DNA double-strand breaks mediate RNA-driven phase separation of damage-response factors. Nat. Cell Biol. 21:1286–1299.

28. Francia, S., M. Cabrini, V. Matti, A. Oldani, and F. d’Adda di Fagagna. 2016. DICER, DROSHA and DNA damage response RNAs are necessary for the secondary recruitment of DNA damage response factors. J. Cell Sci. 129:1468–1476.

29. Durut, N., and O. Mittelsten Scheid. 2019. The Role of Noncoding RNAs in Double-Strand Break Repair. Front. Plant Sci. 10:1–13.

30. Schreiber, V., F. Dantzer, J.-C. Ame, and G. de Murcia. 2006. Poly(ADP-ribose): novel functions for an old molecule. Nat. Rev. Mol. Cell Biol. 7:517–528.

31. Gupte, R., Z. Liu, and W.L. Kraus. 2017. PARPs and ADP-ribosylation: recent advances linking molecular functions to biological outcomes. Genes Dev. 31:101–126.

32. Fay, M.M., and P.J. Anderson. 2018. The Role of RNA in Biological Phase Separations. J. Mol. Biol. 430:4685–4701.

33. Singatulina, A.S., L. Hamon, M. V. Sukhanova, B. Desforges, V. Joshi, A. Bouhss, O.I. Lavrik, and D. Pastré. 2019. PARP-1 Activation Directs FUS to DNA Damage Sites to Form PARG-Reversible Compartments Enriched in Damaged DNA. Cell Rep. 27:1809–1821.e5.

34. Polo, L.M., Y. Xu, P. Hornyak, F. Garces, Z. Zeng, R. Hailstone, S.J. Matthews, K.W. Caldecott, A.W. Oliver, and L.H. Pearl. 2019. Efficient Single-Strand Break Repair Requires Binding to Both Poly(ADP-Ribose) and DNA by the Central BRCT Domain of XRCC1. Cell Rep. 26:573–581.e5.

35. Leung, A.K.L. 2020. Poly(ADP-ribose): A Dynamic Trigger for Biomolecular Condensate Formation. Trends Cell Biol. 30:370–383.

36. Abbotts, R., and D.M. Wilson. 2017. Coordination of DNA single strand break repair. Free Radic. Biol. Med. 107:228–244.

37. Sukhanova, M. V. 2005. Human base excision repair enzymes apurinic/apyrimidinic endonuclease1 (APE1), DNA polymerase and poly(ADP-ribose) polymerase 1: interplay between strand-displacement DNA synthesis and proofreading exonuclease activity. Nucleic Acids Res. 33:1222–1229.

38. Dantzer, F., G. de la Rubia, J. Ménissier-de Murcia, Z. Hostomsky, G. de Murcia, and V. Schreiber. 2000. Base Excision Repair Is Impaired in Mammalian Cells Lacking Poly(ADP-ribose) Polymerase-1. Biochemistry. 39:7559–7569.

39. Dutertre, M., S. Lambert, A. Carreira, M. Amor-Guéret, and S. Vagner. 2014. DNA damage: RNA-binding proteins protect from near and far. Trends Biochem. Sci. 39:141–149.

40. Yuan, M., C.G. Eberhart, and M. Kai. 2014. RNA binding protein RBM14 promotes radio-resistance in glioblastoma by regulating DNA repair and cell differentiation. Oncotarget. 5:2820–2826.

41. Simon, N.E., M. Yuan, and M. Kai. 2017. RNA-binding protein RBM14 regulates dissociation and association of non-homologous end joining proteins. Cell Cycle. 16:1175–1180.

42. Alfano, L., A. Caporaso, A. Altieri, M. Dell’Aquila, C. Landi, L. Bini, F. Pentimalli, and A. Giordano. 2019. Depletion of the RNA binding protein HNRNPD impairs homologous recombination by inhibiting DNA-end resection and inducing R-loop accumulation. Nucleic Acids Res. 47:4068–4085.

43. Taschuk, F., and S. Cherry. 2020. DEAD-Box Helicases: Sensors, Regulators, and Effectors for Antiviral Defense. Viruses. 12:181.

44. Sen, N.D., N. Gupta, S. K Archer, T. Preiss, J.R. Lorsch, and A.G. Hinnebusch. 2019. Functional interplay between DEAD-box RNA helicases Ded1 and Dbp1 in preinitiation complex attachment and scanning on structured mRNAs in vivo. Nucleic Acids Res. 47:8785–8806.

45. Song, C., A. Hotz-Wagenblatt, R. Voit, and I. Grummt. 2017. SIRT7 and the DEAD-box helicase DDX21 cooperate to resolve genomic R loops and safeguard genome stability. Genes Dev. 31:1370–1381.

46. Li, L., D.R. Germain, H.-Y. Poon, M.R. Hildebrandt, E.A. Monckton, D. McDonald, M.J. Hendzel, and R. Godbout. 2016. DEAD Box 1 Facilitates Removal of RNA and Homologous Recombination at DNA Double-Strand Breaks. Mol. Cell. Biol. 36:2794–2810.

47. Nott, T.J., E. Petsalaki, P. Farber, D. Jervis, E. Fussner, A. Plochowietz, T.D. Craggs, D.P. Bazett-Jones, T. Pawson, J.D. Forman-Kay, and A.J. Baldwin. 2015. Phase Transition of a Disordered Nuage Protein Generates Environmentally Responsive Membraneless Organelles. Mol. Cell. 57:936–947.

48. Hondele, M., R. Sachdev, S. Heinrich, J. Wang, P. Vallotton, B.M.A. Fontoura, and K. Weis. 2019. DEAD-box ATPases are global regulators of phase-separated organelles. Nature. 573:144–148.

49. Francia, S., F. Michelini, A. Saxena, D. Tang, M. de Hoon, V. Anelli, M. Mione, P. Carninci, and F. d’Adda di Fagagna. 2012. Site-specific DICER and DROSHA RNA products control the DNA-damage response. Nature. 488:231–235.

50. Antoniali, G., M.C. Malfatti, and G. Tell. 2017. Unveiling the non-repair face of the Base Excision Repair pathway in RNA processing: A missing link between DNA repair and gene expression? DNA Repair (Amst). 56:65–74.

51. Vascotto, C., D. Fantini, M. Romanello, L. Cesaratto, M. Deganuto, A. Leonardi, J.P. Radicella, M.R. Kelley, C. D’Ambrosio, A. Scaloni, F. Quadrifoglio, and G. Tell. 2009. APE1/Ref-1 Interacts with NPM1 within Nucleoli and Plays a Role in the rRNA Quality Control Process. Mol. Cell. Biol. 29:1834–1854.

52. Tell, G., D.M. Wilson, and C.H. Lee. 2010. Intrusion of a DNA Repair Protein in the RNome World: Is This the Beginning of a New Era? Mol. Cell. Biol. 30:366–371.

53. Ayyildiz, D., G. Antoniali, C. D’Ambrosio, G. Mangiapane, E. Dalla, A. Scaloni, G. Tell, and S. Piazza. 2020. Architecture of The Human Ape1 Interactome Defines Novel Cancers Signatures. Sci. Rep. 10:1–19.

54. Piovesan, D., F. Tabaro, L. Paladin, M. Necci, I. Mičetić, C. Camilloni, N. Davey, Z. Dosztányi, B. Mészáros, A.M. Monzon, G. Parisi, E. Schad, P. Sormanni, P. Tompa, M. Vendruscolo, W.F. Vranken, and S.C.E. Tosatto. 2018. MobiDB 3.0: More annotations for intrinsic disorder, conformational diversity and interactions in proteins. Nucleic Acids Res. 46:D471–D476.

55. Mészáros, B., G. Erdős, B. Szabó, É. Schád, Á. Tantos, R. Abukhairan, T. Horváth, N. Murvai, O.P. Kovács, M. Kovács, S.C.E. Tosatto, P. Tompa, Z. Dosztányi, and R. Pancsa. 2019. PhaSePro: the database of proteins driving liquid-liquid phase separation. Nucleic Acids Res. 48:D360–D367.

56. You, K., Q. Huang, C. Yu, B. Shen, C. Sevilla, M. Shi, H. Hermjakob, Y. Chen, and T. Li. 2020. PhaSepDB: a database of liquid-liquid phase separation related proteins. Nucleic Acids Res. 48:D354–D359.

57. Subramanian, A., P. Tamayo, V.K. Mootha, S. Mukherjee, B.L. Ebert, M.A. Gillette, A. Paulovich, S.L. Pomeroy, T.R. Golub, E.S. Lander, and J.P. Mesirov. 2005. Gene set enrichment analysis: A knowledge-based approach for interpreting genome-wide expression profiles. Proc. Natl. Acad. Sci. U. S. A. 102:15545–15550.

58. Bindea, G., B. Mlecnik, H. Hackl, P. Charoentong, M. Tosolini, A. Kirilovsky, W.-H. Fridman, F. Pagès, Z. Trajanoski, and J. Galon. 2009. ClueGO: a Cytoscape plug-in to decipher functionally grouped gene ontology and pathway annotation networks. Bioinformatics. 25:1091–1093.

59. Shannon, P., A. Markiel, O. Ozier, N.S. Baliga, J.T. Wang, D. Ramage, N. Amin, B. Schwikowki, and T. Ideker. 2003. Cytoscape: A Software Environment for Integrated Models of Biomolecular Interaction Networks. Genome Res. 13:2498–2504.

60. Wegmann, S., B. Eftekharzadeh, K. Tepper, K.M. Zoltowska, R.E. Bennett, S. Dujardin, P.R. Laskowski, D. MacKenzie, T. Kamath, C. Commins, C. Vanderburg, A.D. Roe, Z. Fan, A.M. Molliex, A. Hernandez-Vega, D. Muller, A.A. Hyman, E. Mandelkow, J.P. Taylor, and B.T. Hyman. 2018. Tau protein liquid–liquid phase separation can initiate tau aggregation. EMBO J. 37:1–21.

61. Aguzzi, A., and M. Altmeyer. 2016. Phase Separation: Linking Cellular Compartmentalization to Disease. Trends Cell Biol. 26:547–558.

62. Liberzon, A., A. Subramanian, R. Pinchback, H. Thorvaldsdóttir, P. Tamayo, and J.P. Mesirov. 2011. Molecular signatures database (MSigDB) 3.0. Bioinformatics. 27:1739–1740.

63. Liberzon, A., C. Birger, H. Thorvaldsdóttir, M. Ghandi, J.P. Mesirov, and P. Tamayo. 2015. The Molecular Signatures Database Hallmark Gene Set Collection. Cell Syst. 1:417–425.

64. Bader, A.S., B.R. Hawley, A. Wilczynska, and M. Bushell. 2020. The roles of RNA in DNA double-strand break repair. Br. J. Cancer. 122:613–623.

65. Alberti, S., S. Saha, J.B. Woodruff, T.M. Franzmann, J. Wang, and A.A. Hyman. 2018. A User’s Guide for Phase Separation Assays with Purified Proteins. J. Mol. Biol. 430:4806–4820.

66. Conicella, A.E., G.H. Zerze, J. Mittal, and N.L. Fawzi. 2016. ALS Mutations Disrupt Phase Separation Mediated by α-Helical Structure in the TDP-43 Low-Complexity C-Terminal Domain. Structure. 24:1537–1549.

67. Jain, A., and R.D. Vale. 2017. RNA phase transitions in repeat expansion disorders. Nature. 546:243–247.

68. Elbaum-Garfinkle, S. 2019. Matter over mind: Liquid phase separation and neurodegeneration. J. Biol. Chem. 294:7160–7168.

